# Uncovering complex disease subtypes by integrating clinical data and imputed transcriptome from genome-wide association studies: Applications in psychiatry and cardiovascular medicine

**DOI:** 10.1101/595488

**Authors:** Liangying Yin, Carlos K.L. Chau, Pak-Chung Sham, Hon-Cheong So

**Affiliations:** School of Biomedical Sciences, Faculty of Medicine, The Chinese University of Hong Kong; Centre for Genomic Sciences, University of Hong Kong; Department of Psychiatry, University of Hong Kong; State Key Laboratory for Cognitive and Brain Sciences, University of Hong Kong; KIZ-CUHK Joint Laboratory of Bioresources and Molecular Research of Common Diseases, Kunming Zoology Institute of Zoology and The Chinese University of Hong Kong; Department of Psychiatry, The Chinese University of Hong Kong; Margaret K.L. Cheung Research Centre for Management of Parkinsonism, The Chinese University of Hong Kong; Shenzhen Research Institute, The Chinese University of Hong Kong

## Abstract

Classifying patients into clinically and biologically homogenous subgroups will facilitate the understanding of disease pathophysiology and development of more targeted prevention and intervention strategies. Traditionally, disease subtyping is based on clinical characteristics alone, however disease subtypes identified by such an approach may not conform exactly to the underlying biological mechanisms. Very few studies have integrated *genomic profiles* (such as those from GWAS) with clinical symptoms for disease subtyping.

In this study, we proposed a novel analytic framework capable of finding subgroups of complex diseases by leveraging both GWAS-predicted gene expression levels and clinical data by a multi-view bicluster analysis. This approach connects SNPs to genes via their effects on expression, hence the analysis is more biologically relevant and interpretable than a pure SNP-based analysis. Transcriptome of different tissues can also be readily modelled. We also proposed various new evaluation or validation metrics, such as a newly modified ‘prediction strength’ measure to assess generalization of clustering performance. The proposed framework was applied to derive subtypes for schizophrenia, and to stratify subjects into different levels of cardiometabolic risks.

Our framework was able to subtype schizophrenia patients with diverse prognosis and treatment response. We also applied the framework to the Northern Finland Cohort (NFBC) 1966 dataset, and identified high- and low cardiometabolic risk subgroups in a gender-stratified analysis. Our results suggest a more data-driven and biologically-informed approach to defining metabolic syndrome. The prediction strength was over 80%, suggesting that the cluster model generalizes well to new datasets. Moreover, we found that the genes ‘blindly’ selected by the cluster algorithm are significantly enriched for known susceptibility genes discovered in GWAS of schizophrenia and cardiovascular diseases, providing further support to the validity of our approach. The proposed framework may be applied to any complex diseases, and opens up a new approach to patient stratification.

## Introduction

Accurate classification of complex diseases such as psychiatric and cardiometabolic disorders into clinically and biologically homogenous subtypes could facilitate the understanding of disease pathophysiology and development of more targeted interventions^1^. Traditionally, disease subtyping are based on clinical characteristics alone, however disease subtypes identified by such an approach may not conform exactly to the underlying biological mechanisms. For example, the same disease symptom may be caused by different mechanisms in different patients. Patients with similar clinical presentations can also have varying response to treatment. On the other hand, last decade has witnessed the remarkable success of genome-wide association studies (GWAS) in identifying susceptibility loci for complex diseases^2^. In addition to yielding mechanistic insights into various disorders, GWAS data may also be useful in a more directly translational context. For example, there has been increasing interest to apply GWAS data for risk prediction^3^ and drug discovery or repurposing^4^. However, despite >3000 GWAS being performed (https://www.ebi.ac.uk/gwas/), another potential translational application has been largely ignored: could genomic information from GWAS help to improve *patient stratification or disease subtyping*? As argued above, subtyping by disease symptoms or characteristics alone has its limitations, which may be improved upon by the combination of both clinical and genomic information.

Very few works have studied on how genomic data from GWAS may reveal complex disease subtypes. Arnedo et al. investigated genetic architecture of schizophrenia by independently identifying SNP- and phenotype ‘sets’ and studying their inter-relationships ^5^. However, there are other limitations, for example the number of subgroups are allowed to vary in a very wide range (up to ~90). Besides, there are potential problems with significance testing (For more details, please see: http://genomesunzipped.org/2014/09/eight-types-of-schizophrenia-not-so-fast.php). Cleynen et al.^6^ performed disease subtyping on Crohn’s disease based on 46 single nucleotide polymorphisms (SNPs) extracted from a GWAS and found modest differences in clinical variables among the subgroups.

In a more recent work^8^, we have presented the first application of whole-genome SNP data and clinical variables for subtyping a complex disease. We studied schizophrenia (SCZ), a highly heterogeneous psychiatric disorder. We found that the identified subgroups were indeed different with respect to treatment response and other outcome variables, providing support to the use of genetic data in disease subtyping. However, there are several important limitations regarding this SNP-based approach to subtyping. Firstly, the functional roles of many SNPs identified in GWAS remain unknown^9^. Previous studies reported that the majority (up to ~88%)^10^ of GWAS tag SNPs lie in intergenic or intronic regions. While the cluster algorithm could identify a subset of SNPs that characterize each cluster, the results could be difficult to interpret, as most SNPs do not have clear functional implications, and many are intronic or intergenic. In addition, as some SNPs cannot be easily mapped to genes, subsequent gene-based analysis (e.g. on pathway enrichment) may be suboptimal. Secondly, the dimension of SNPs is extremely high and could reach >10 million with imputation. While an alternative approach is to perform pre-screening for a subset of more promising variants before cluster analysis, the choice of the significance threshold for SNP inclusion is often arbitrary. In addition, our previous work also showed inferior performance of a pre-screening approach compared to modelling all SNPs ^8^. Also, it has been argued recently that a very large number of genetic variants, or even the majority of the genome, may be associated with complex diseases^11^. Hence restricting analysis to a subset of highly significant SNPs may miss the contribution of many true disease variants. Nevertheless, there is a major problem in analysing all SNPs: when the dimension of features (e.g. SNPs) is very high, the computational burden of cluster analysis will likely become heavy, especially with large sample sizes. The SNP-based analysis may then become impractical due to slow computational speed and heavy memory requirements.

In this study, we propose a novel analytic framework capable of finding subgroups of complex diseases. One of the key innovations is to leverage GWAS-predicted gene expression levels instead of raw SNP data. Estimating gene expression from genotype has become an active area of research, thanks to increasing eQTL resources such as GTEx and others^12,13^. In our work, genomic data are combined with clinical phenotypes, and *both* types of data are utilized for disease subtyping via an (unsupervised) machine learning approach known as ‘multi-view clustering’. The overall aim is to classify patients into meaningful subgroups with clinical and biological significance. The new gene-based approach is considerably faster and much less memory-demanding than the SNP-based approach; more importantly, the approach connects the functional impact of the SNPs to genes via their effect on expression for subsequent cluster analysis. Changes in expression levels may be closer to the underlying pathophysiology of diseases, and the results from the analysis are easier to interpret. Another important advantage is that we can impute expression levels in *different tissues* easily, while it is impossible to consider tissue relevance if we model the SNPs directly.

In oncology, finding molecular subtypes of cancer characterized by different expression and other omics profiles has been an active area of research, and showed great promise for translating into more targeted intervention and prognostic strategies for patients. One of the reasons for more active subtyping studies in oncology might be due to the availability of relevant tissues from surgical specimens; omics profiles can then be measured, often in samples without prior drug treatment (e.g. TCGA samples are free from neoadjuvant treatment). For other complex diseases, such as psychiatric disorders, access to relevant tissues is usually invasive and costly (or requires post-mortem samples), and expression data are often confounded by medications taken. Our proposed approach using GWAS-imputed transcriptome data avoids these issues as imputation can be based on external reference data, hence expression levels in different tissues can be easily imputed without invasive procedures. Also the results are not affected by drug use, non-pharmacological interventions or other environmental confounders, as our imputation is based on (germline) genetic variants.

We evaluated the feasibility and validity of our proposed approach on two different categories of complex diseases, namely psychiatric and cardiometabolic disorders/traits. We presented an analytic framework for disease subtyping, and also proposed several new validation strategies to check the validity of the clustering algorithm and derived disease subtypes. Our presented analytic framework is general and may be applied to any complex diseases. Our results indicated that the proposed approach has the capability to stratify patients into meaningful subgroups with clinical and biological relevance.

## Method

The main purpose of this study is to present a novel analytic framework to discover disease subtypes through incorporating GWAS-predicted expression levels and clinical traits. We employed a multi-view clustering method, which is capable of uncovering disease heterogeneity across different data views of patients (clinical and genetic). A schematic diagram of our proposed approach is shown in Fig.1. Our method includes three steps, i.e., data ‘imputation’, disease subtypes discovery and validation of discovered subtypes (through internal and external validation approaches). We shall describe each of the steps in greater detail below.

**Fig. 1.**
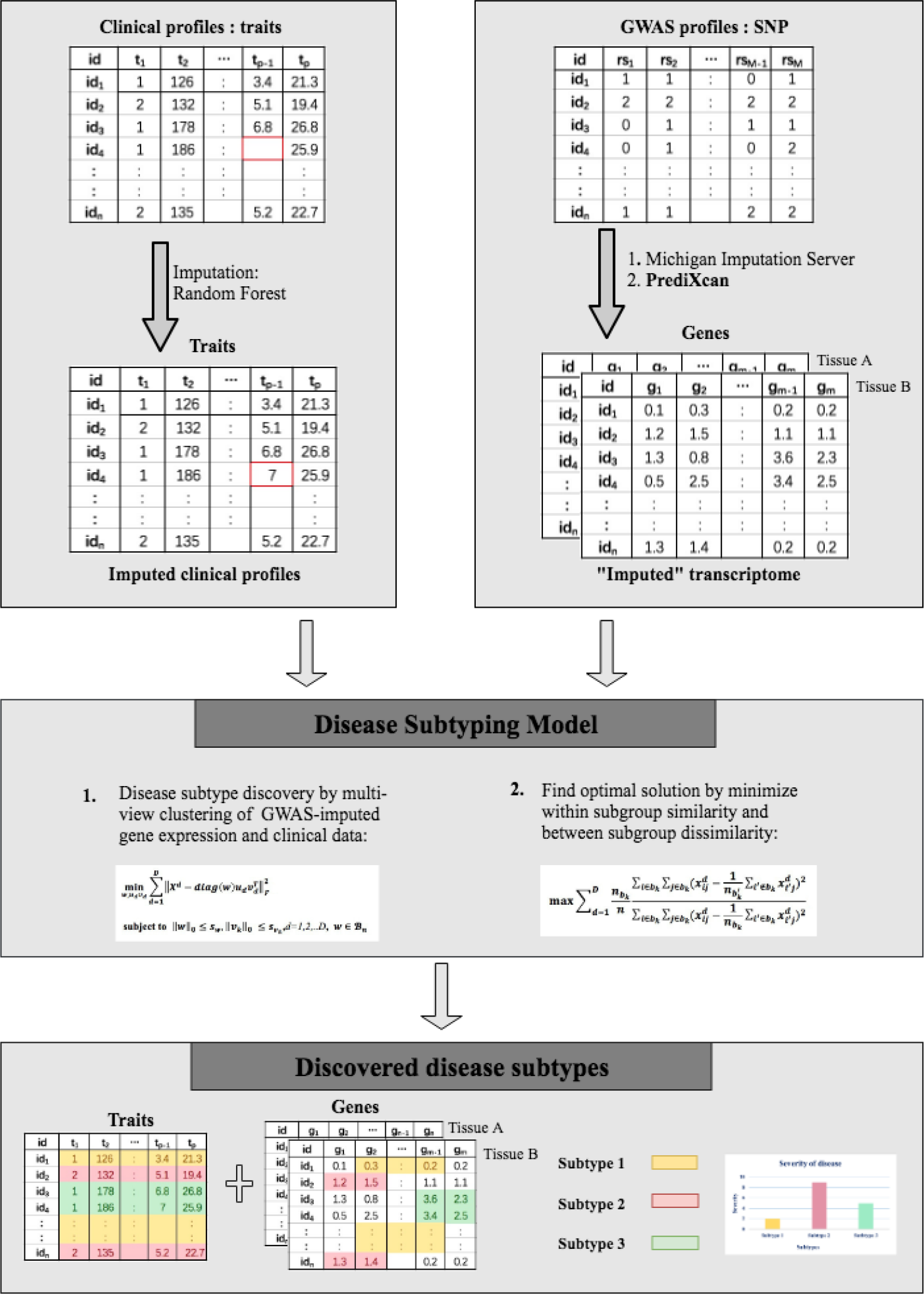
Outline of proposed method for disease subtypes discovery

In brief, our method includes three steps, i.e., data ‘imputation’, disease subtypes discovery and validation of discovered subtypes. 1) data ‘imputation’: For clinical traits, we apply R package ‘missForest’ which employs a random forest approach for data imputation; Then, we employed ‘PrediXcan’ to map our SNPs to genes with estimated gene expression levels. 2) disease subtypes discovery: we make use of a multi-view sparse clustering method to uncover the underlying subtypes of complex disease by utilizing both GWAS-imputed gene expression levels and clinical data. 3) Validation of discovered subtypes: both internal and external validation were conducted.

### Data imputation and estimation of expression levels from GWAS

As missing data are not allowed for clustering analysis, imputation was firstly performed. For clinical traits, we apply the R package ‘missForest’ for data imputation which employs a random forest algorithm for imputation ^14^. For the GWAS dataset, we need to impute expression levels, which is best conducted on a full set of genetic variants instead of SNPs on the genotyping panel only. We therefore performed variant-level imputation by the program ‘Minimac’ using the University of Michigan Imputation Server and 1000 Genomes Phase 3 v5 as the reference panel^15^. SNPs with INFO score> 0.3 were kept. We then employed ‘PrediXcan’ to impute expression levels from the imputed genotype data. For details of the algorithm, please refer to original paper ^16^. Briefly, the algorithm first produces prediction models for expression levels from an external reference dataset (such as GTEx) which contains both genotype and expression data. An elastic net regression model is used by default. Then, the prediction model can be applied to new genotype data to ‘impute’ expression. Expression in different tissues can be estimated as long as the reference dataset includes such data.

### Disease subtype discovery

#### Multi-view sparse biclustering

Supposing *X*^*d*^ is a *n* × *m*_*d*_ data matrix from clinical or genetic view of patients, where *n* is the sample size, *d* denotes the index of ‘view’ to be modelled and *m*_*d*_ is the number of features in the *d*^th^ view. For example, if one models clinical and GWAS-predicted expression in one tissue, there will be two views. It is possible to extend the approach to more than 2 views, for example based on expression in different tissues or using other (preferably gene-based) ‘omics’ profiles. Subgroup of patients can be simultaneously derived by performing a *sparse rank one approximation* on the original matrices *X*^*d*^(*d* = 1,2,.. *D*, indicating data matrices from different *views* that characterize the same set of patients), i.e.,

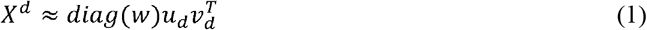

where *w* is a binary vector of size *n*, serving as a common factor that force different views of data to agree on the same grouping of patients. *diag(w)* is a diagonal matrix of size *n* × *n* with diagonal entries equal to *w*. *u*_*d*_ of size *n* and *v*_*d*_ of size *m*_*d*_ are the rank-one approximations of *X*^*d*^ respectively. Rows in *X*^*d*^ corresponding to the non-zero entries of *diag(w)* form the row subgroups, and columns in *v*_*d*_ form the column subgroups (a.k.a., sub-feature groups) in different views. Subgroups of patients based on different views of data can be derived by solving the following optimization problem:

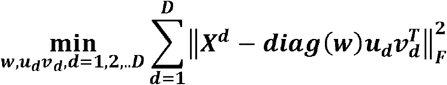

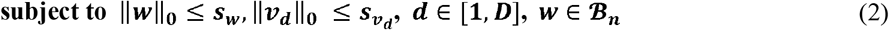

where ***s***_***w***_ and ***s***_*v*_*d*__’s are hyper-parameters that need to be predetermined to enforce sparsity of *w* and *v*_*d*_’s, i.e., the number of patients *n*_*b*_*k*__ and number of selected features 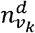 in each subgroup of the corresponding data view. *D* is the number of data views incorporated for clustering and *B*_*n*_ is the set that contains all possible binary vectors of length n. To obtain subsequent subgroups, we need to firstly update the data matrices by excluding previously identified patients, then solve Eq. (2). For details of optimization of the objective function, please refer to the original paper ^17^.

The presented approach is capable of selecting features during the clustering process, however we need to predetermine the number of selected features in each data view. For data matrix from clinical view of patients, all features were preserved for disease subgroup discovery. As for data matrices from genetic views of patients, based on suggestions from the original paper we employed principal component analysis (PCA) to determine the number of selected features 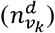 in each view. As recommended by the authors, we set 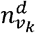 to be the number where the accumulated variance in PCA of ***X***^***d***^ (genetic view) was over 90%.

As for the number of possible disease subgroups, we considered a range of 2 to 6 subgroups. We need to determine the number of patients in each disease subgroup in each clustering trial. Here we set the smallest number of patients [*min*(*n*_*b*_*k*__)] in each subgroup to 20. For a given number of subgroups *k*, we firstly set *n*_*b*_*k*__ to a value roughly equals to *n*/*k*, then we experimented with all the combinations by adding or subtracting *min*(*n*_*b*_*k*__) in each subgroup. For example, suppose we have 400 patients and *k* = 2, then *n*_*b*_1__ and *n*_*b*_2__ would be firstly set to 200, subsequently we would experiment with *n*_*b*_1__ = 200 + 20\**t* & *n*_*b*_2__ = 200 − 20\**t* (*t* =1,2,..9).

### Finding optimal biclusters by comparing between- and within-bicluster distances

In order to find the optimal solution for disease subgroups, we proposed a new algorithm to evaluate the identified biclusters in given datasets. One of the most commonly employed index for bicluster performance is the mean squared residue (MSR)^18^, which assesses the homogeneity within each bicluster. However, the index does not maximize the *heterogeneity* between different biclusters. For well-separated biclusters, patients within the same subgroup should be highly homogeneous while patients belong to different subgroups should be highly heterogeneous. In this regard, finding well-separated biclusters (subgroups of patients) is equivalent to finding multi-view clustering results that maximize the sum of ratios of between bicluster distance and within bicluster distance 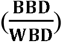 over all data views and all biclusters. Hence we came up with the following index

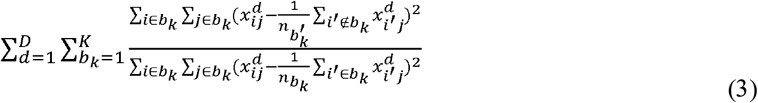

where *i* and *j* are the index of patients and features in the derived subgroup *b*_*k*_, 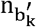 is the number of patients in the given datasets that not belong to subgroup *b*_*k*_ while *n*_*b*_*k*__ is the number of patients in subgroup *b*_*k*_.

We note that if a bicluster is of smaller sample size, the 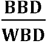 of this bicluster tends to be larger, as it is easier to achieve a smaller within-bicluster variance. To remedy this potential bias, we weighted the 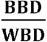 of each bicluster based on their sample size proportionally, i.e.,

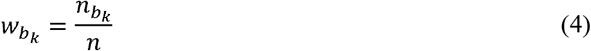

hence imposing a penalty for smaller biclusters. We identify the best solution by finding the bicluster configuration that maximizes the following objective function:

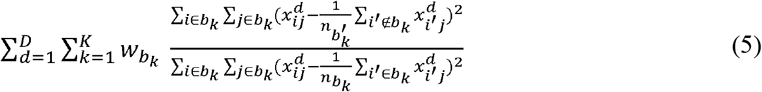

### Evaluation of discovered subgroups

We employed two approaches to validate the discovered patient subgroups. When external data on disease outcome (that are *not* involved in the clustering process) is available, one may validate the derived patient subgroups by finding differences in prognosis across subgroups. On the other hand, if such data is not available, one may employ other internal validation methods. In this study, we presented an approach that involved splitting the sample into ‘training’ and ‘testing’ sets, and evaluated whether the patient subtyping model derived from the training set ‘predicts’ the actual subgroups derived from the testing set alone. The methods are detailed below.

#### External validation

To assess the validity of the discovered disease subgroups, we compared the identified disease subgroups to a number of outcome-related variables that were *not* used for clustering. We performed regression analysis to evaluate the differences among discovered disease subgroups. Ordinal regression was applied for ordered responses of >= 3 groups. We employed the Benjamini-Hochberg false discovery rate approach (FDR) to control for multiple testing. FDR controls the expected proportion of false positive results among those declared significant.

#### Internal validation

In case there are no available outcomes for external validation, internal validation approach is required to assess the quality of the discovered disease subgroups. Here we proposed using a modified version of “prediction strength” (PS) for validation. PS is a widely used validation metric proposed by Tibshirani and Walther to measure the quality of (single-view) *k*-means clustering results. Here we developed a new modified version of this metric for use in biclustering analysis, which has not been reported before. Details on the original PS algorithm can be found in the original paper^19^. Briefly, PS can be conceptualized as an extension of cross-validation used in supervised learning problems. The data is randomly split into a train-set and a test-set, and clustering is performed on each set separately. The clustering model from the train-set is then applied onto the test-set; this is usually done by assigning each observation in the test-set to the nearest cluster centroid derived from the train-set. One can then compute how well the co-memberships based on the ‘predicted’ clusters matches with the co-memberships derived from actually performing another cluster analysis in the test-set. The prediction strength therefore enables one to assess how well the cluster model can be generalized to new datasets, analogous to examining the predictive performance of a supervised classification model in a new dataset.

As single-view *k*-means clustering is different from the multi-view sparse biclustering that we employed, we proposed a new metric to compute PS. As described above, the algorithm involves assigning cluster labels to test-set observations according to the clustering model derived from the train-set. To perform this step, we calculate the distance between each *test*-set observation and derived *train*-set cluster centres. However, unlike in ordinary clustering where all the features are employed, here only a subset of the features are selected in each bicluster, and the selected features could vary in different biclusters. If we just compute the distances between each observation and bicluster centers, the comparison is not fair as the features used for distance calculation are different for the *k* biclusters.

We therefore propose a new approach for distance computation, enabling the comparison to be done on the same set of features. Say for an example, three biclusters have been derived, and the selected features sets were *A*, *B* and *C* respectively. For a new observation *x*_new_, we first compute the distance of *x*_new_ to the center of bicluster 1, considering feature-set *A* only; then again based on feature-set *A*, we compute the distance of *x*_new_ to the center of a ‘combined’ bicluster formed by subjects belonging to biclusters 2 and 3. We take the ratio of these two distances as the new measure of proximity to bicluster 1. The procedure is repeated for bicluster 1, 2…*k*, and each test observation is assigned to the bicluster with the lowest ratio of distances as derived above. In equation form, the new proximity measure of each test observation (*x*_*i*_) to a bicluster (*b*_*k*_) can be expressed as

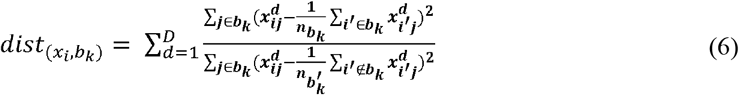

where *i* is the index of patients in test set, *j* is the index of features, *b*_*k*_ is the derived bicluster in training set, *n*_*b*_*k*__ is the number of patients in the bicluster *b*_*k*_ while 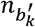 is the number of patients who do not belong to bicluster *b*_*k*_. The process of distance computation is illustrated in Figure 2. Each patient is assigned to its nearest bicluster, i.e., min *k*{*d*_(*t*_*i*_,*b*_*k*_)_|*k* = 1,2,…, *K*}. The prediction strength of a clustering process can be calculated by

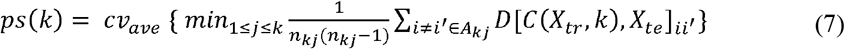

Where *C*(*X*_*tr*_,*k*) denote the clustering operation on training set, *D*[*C*(*X*_*tr*_,*k*),*X*_*te*_]_*ii′*_ is the co-membership matrix with *D*[*C*(*X*_*tr*_,*k*),*X*_*te*_]_*ii′*_ = 1 if patients *i* and *i′* fall into the same bicluster and *D*[*C*(*X*_*tr*_,*k*),*X*_*te*_]_*ii′*_ =0 otherwise. *cv*_ave_ refers to taking the average across all cross-validation folds. In this study, following the original study, we randomly split the sample into 2 halves and performed 2-fold CV 3 times. According to Tibshirani and Walther *ps*(*k*) ≥ 0.8 suggests well-separated biclusters.

**Fig. 2.**
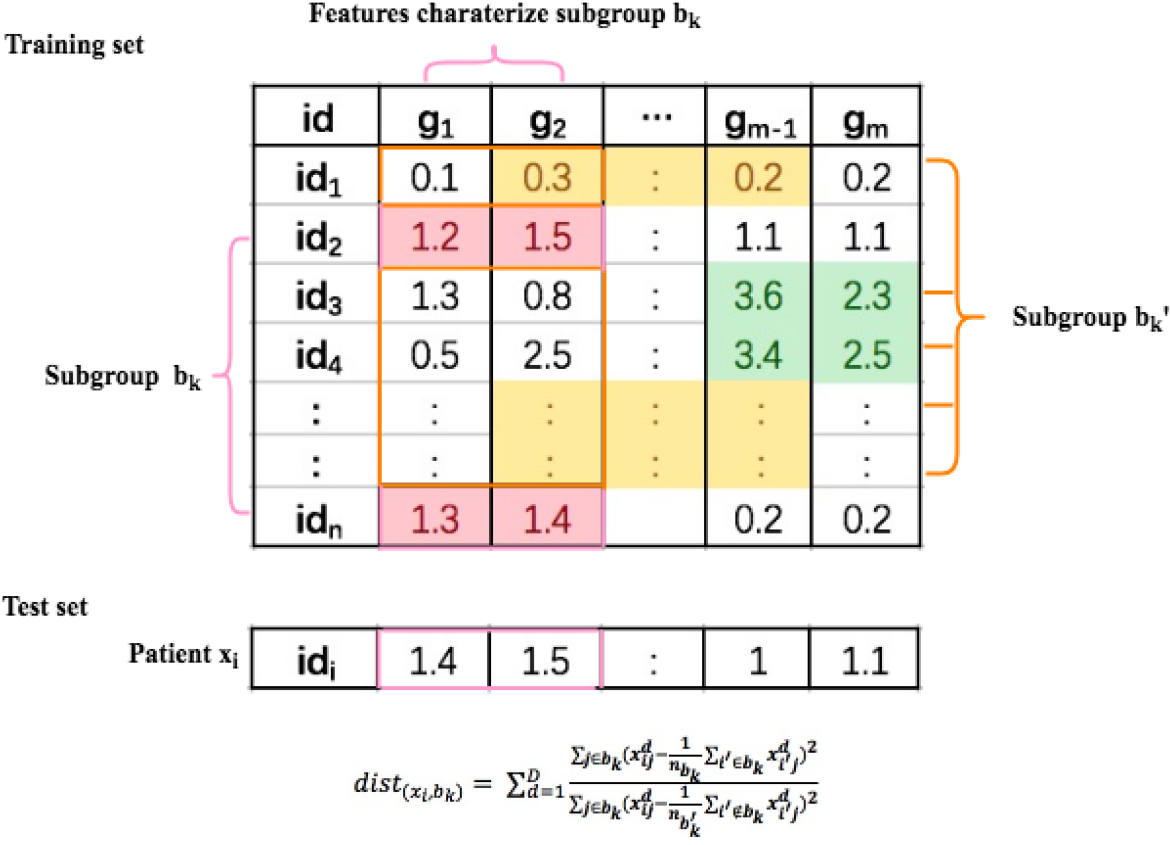
Illustration of distance calculation for prediction strength

#### To confirm the presence of cluster structure in our data

To verify that the discovered clusters are “really there” instead of the results of natural sampling variation, we employed the R package “sigClust” to test for the presence of cluster structure in our data. We used the settings suggested by the authors. Details on this algorithm are described elsewhere ^20^.

#### Selected genes and pathway analysis

We extracted the genes selected in the clustering process to figure out which genes contribute to the subtyping of patients. We also conducted pathway analysis to further explore the pathophysiology in each When the number of selected genes is relatively small, one may employ an over-representation analysis based on hypergeometric tests. Alternatively, the vector *v*_*k*_ can also be considered as a measure of the weight of different features, and such information can be incorporated into certain pathway analysis algorithms such as GSEA (Gene-set Enrichment Analysis)^21^. We employed the latter approach for pathway analysis in the Northern Finland Birth Cohort sample (see below), as the number of selected genes was relatively large. Over-representation analysis was conducted in “ConsensusPathDB” while GSEA was conducted using WebGastalt ^22,23^. We also performed a “tissue specificity” analysis in FUMA by examining whether selected genes were differentially expressed genes in a particular tissue ^24^.

### Application to real data

#### Subtyping schizophrenia

We applied the proposed framework to 387 schizophrenia (SCZ) patients with clinical, neurocognitive and genetic profiles collected in Hong Kong. Schizophrenia is a psychiatric disorder in which patients are highly heterogeneous with respect to many aspects such as clinical symptoms, prognosis, treatment outcome and probably in the underlying pathogenesis. Same as other psychiatric disorders, the current diagnostic criteria for SCZ relies on clinical symptoms only. Characterization of SCZ patients into more biologically and clinically homogenous subtypes will be an important step towards precision psychiatry. Details of subject recruitment and profile assessment can be found elsewhere. ^25^ Briefly, all subjects met the DSM-IV diagnostic criteria for SCZ. They were all recruited from Hong Kong and were Han Chinese. Clinical characteristics such as course of illness, positive and negative symptom scores, treatment response, history of self-harm and aggression were recorded by trained psychiatrists. Several neurocognitive tests were also performed such as verbal fluency, Stroop test, soft neurological signs and intelligence. All subjects were genotyped by the Illumina Human610-Quad BeadChip and imputation was performed. Standard quality control procedures were conducted following Wong et al^26^.

#### Application to the Northern Finland Birth Cohort (NFBC)

We also applied our proposed approach to the Northern Finland Birth Cohort 1966 (dbGaP Study Accession number phs000276.v2.p1) with a sample size of 4982 (male: 2452, female: 2530). The original study was described in^27^. We performed standard quality control procedures as described earlier^3^. Subjects were genotyped by the Illumina Infinium 370cnvDuo array, and imputation was performed by the Michigan Imputation Server as described above.

In addition to de-identified genome-wide SNP data, a selected list of 13 phenotypes related to cardiovascular disease (CVD) risks including, gender, C-reactive protein, waist-hip ratio (WHR), body mass index (BMI), high-density lipoprotein cholesterol (HDL), low-density lipoprotein cholesterol (LDL), total cholesterol (TC), triglyceride (TG), fasting glucose, fasting insulin, homeostatic model assessment for insulin resistance (HOMA-IR), systolic and diastolic blood pressure (SBP/DBP) were also modeled in our biclustering analysis. Details about phenotype assessment were described elsewhere^25^. All subjects were 31 years of age at the time of assessment. In a recent work, Ongen et al have shown that coronary artery and liver are the top causal tissues for CVD ^28^. In this regard, we selected the imputed gene expression levels of coronary artery and liver along with the clinical profiles of subjects as inputs to our model.

The motivation is to provide a more data-driven and biologically-informed way to stratify subjects into different levels of cardiometabolic risks. At present, individuals are classified as having the ‘metabolic syndrome’ (MetS) if they have several inter-related risk factors (e.g. obesity/central obesity, dyslipidemia, hypertension, hyperglycemia) which leads to increased risk of cardiometabolic diseases. However, the criteria of MetS controversial and different groups^29,30,31^ have proposed different definitions. Also, it is unclear whether MetS truly reflect a subgroup with homogenous pathophysiology^32^. The proposed framework includes genetic factors, which may help identify patient subgroups with more homogenous pathogenic mechanisms; our data-driven approach also reduces the subjectivity in defining cut-offs of metabolic parameters.

## Results

### Application to SCZ patients

We firstly applied our framework to SCZ. We selected imputed gene expression profiles of 10 brain tissues as well as clinical and neurocognitive profile as inputs (*d*=11). Note that we have selected 9 variables which were assessed at baseline or believed to be more stable across the course of illness (e.g. neuro-cognitive measures) as input to the algorithm. The idea is that we wish to subtype the patient at an early stage of illness. On the other hand, another 9 clinical variables related to disease outcome, including history of violence and self-harm, PANSS (The Positive and Negative Syndrome Scale) scores and course of disease, were reserved for validating the differences between derived patient subgroups. These variables were *not* used in the clustering process. Best performance was achieved when patients were categorized into 3 subgroups. Fig.3, Table 1 and Table S1 demonstrate the distributions of clinical and neurocognitive features of patients among the 3 discovered subgroups.

**Fig. 3.**
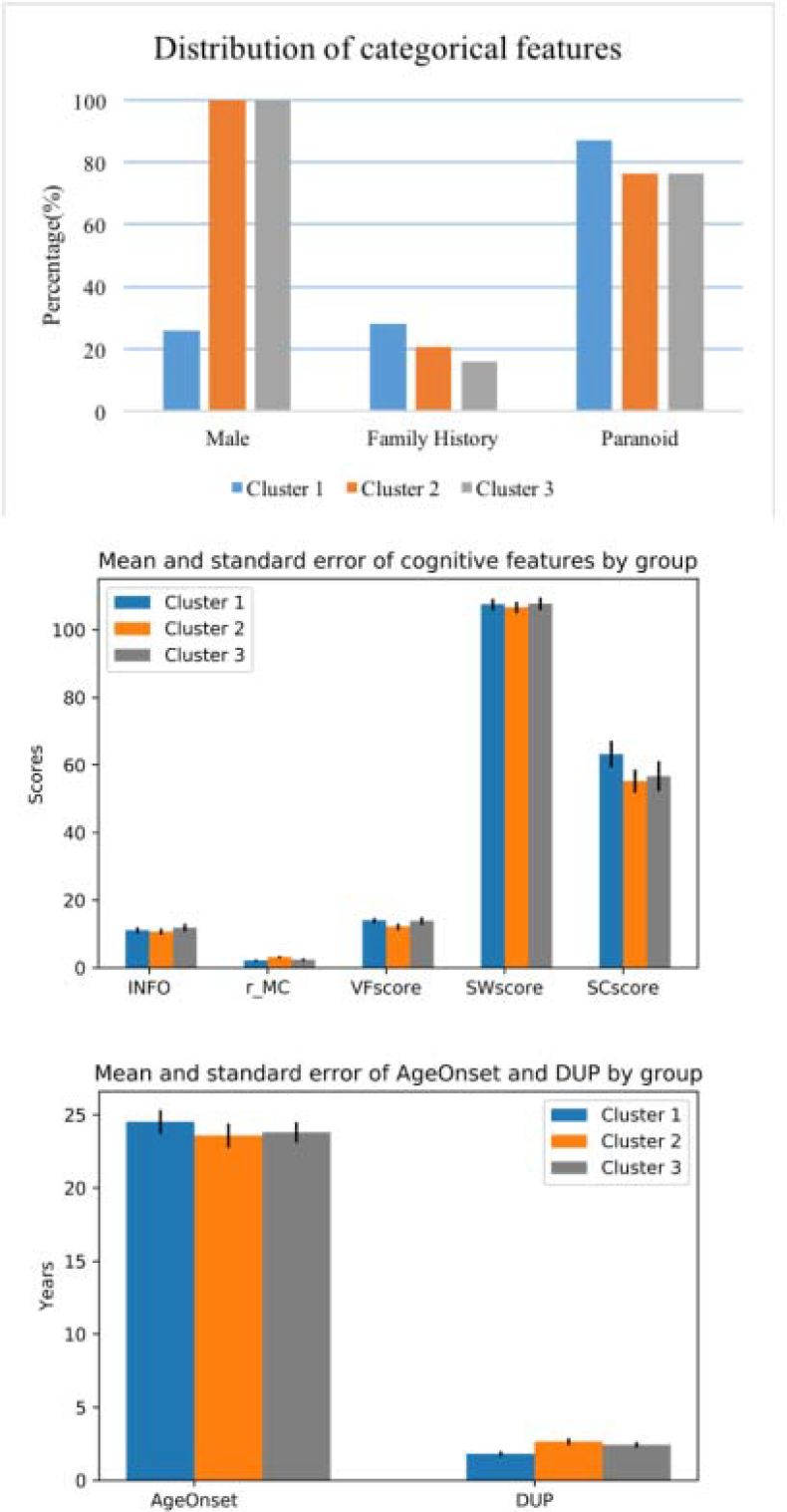
Comparison across input clinical and neurocognitive features by subgroups for SCZ patients

**Table 1.**
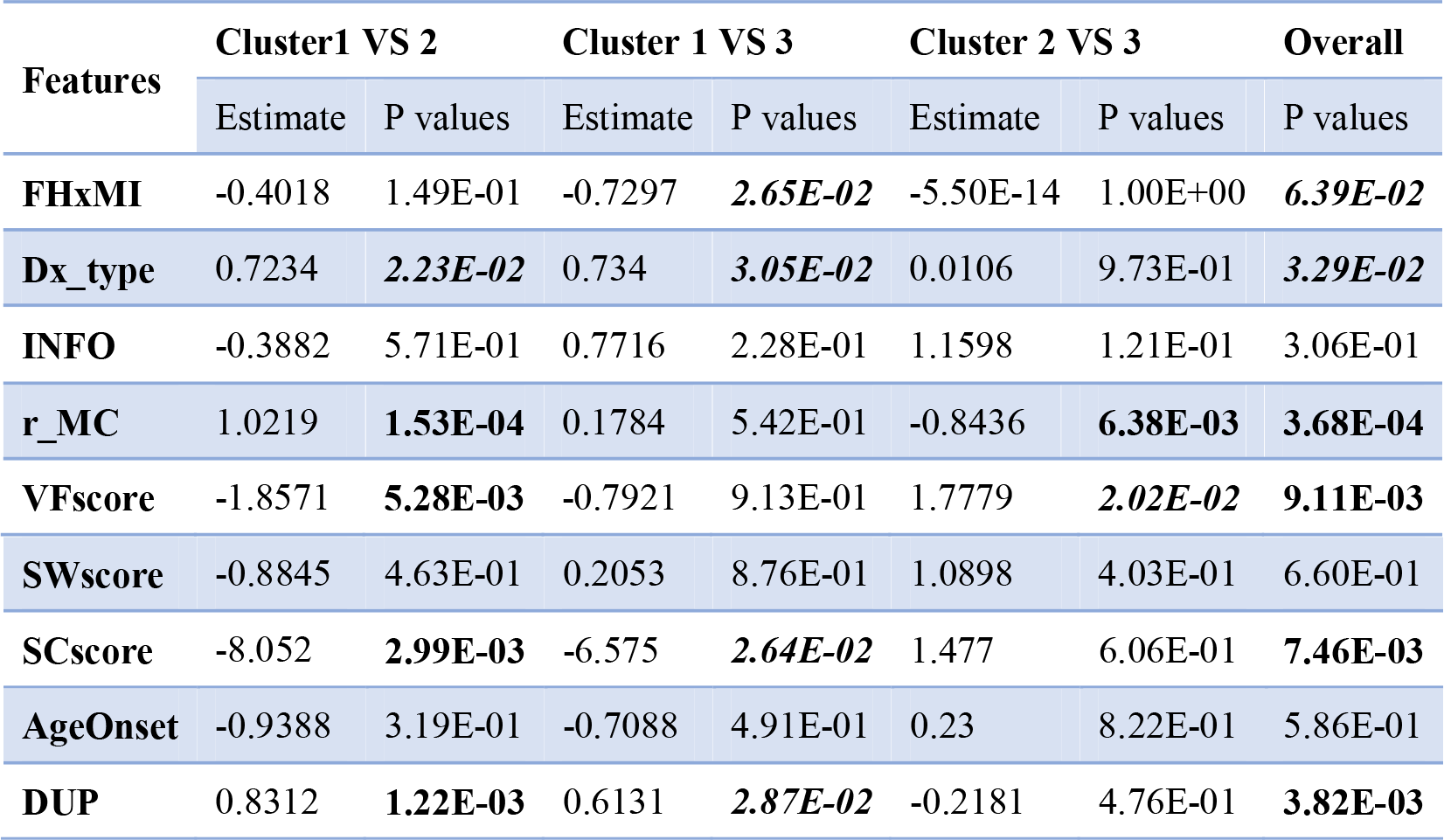
Comparison across input variables for clustering of SCZ patients’ subgroups

As demonstrated in Fig.3, derived patient subgroups showed differences in gender proportions. While 26% of patients were males in subgroup 1, all patients were males in the remaining two subgroups. It is worth noting that gender differences in schizophrenia is well-established^33,34,35^, so imbalance in the male/female proportion across the subgroups are not entirely surprising. Compared with the remaining two subgroups, the first derived subgroup showed a trend towards high proportion of positive family history of mental illness and paranoid schizophrenia. In addition, they had a significant shorter period of untreated psychosis. As for patients in subgroup 2, they had poorer performance on motor coordination and verbal fluency compared to the 1^st^ and 3^rd^ subgroup. Patients in subgroup 3 had intermediate clinical and neurocognitive manifestations.

We then compared the identified subgroups across 9 outcome-related variables. As demonstrated in Fig. 4, Table 2 and Table S2, there existed significant differences among derived subgroups in almost all outcome variables, except for self-harm and aggression subscale of PANSS. In summary, we revealed 3 SCZ subgroups with good, intermediate and poor prognosis. To be more specific, patients in the first subgroup had the lowest tendency for violent behaviours. Besides, they tended to have better treatment response and a more favorable course of disease. For symptom scores, they showed the lowest severity with respect to PANn (negative symptoms), PANg (general psychopathology) and PANtotal (total score). Compared to the first subgroup, the second subgroup exhibited a tendency for poorer treatment response and a continuous course of SCZ. Furthermore, they had the most severe symptoms across almost all subscales of PANSS. Subgroup 3 was intermediate.

**Table 2.** Comparison across outcome-related variables for clustering of SCZ patients’ subgroups

| Features | Cluster1 VS 2 |  | Cluster 1 VS 3 |  | Cluster 2 VS 3 |  | Overall |
| --- | --- | --- | --- | --- | --- | --- | --- |
|  | Estimate | P values | Estimate | P values | Estimate | P values | P values |
| **Violence** | 0.8576 | **1.70E-04** | 1.03601 | **1.88E-05** | -0.1731 | 4.68E-01 | **9.44E-07** |
| **Self-harm** | -0.3438 | 1.71E-01 | -0.4058 | 1.45E-01 | -0.0621 | 8.29E-01 | 1.18E-01 |
| **Treatment response** | 1.2489 | **1.27E-07** | 1.0106 | **6.94E-05** | -0.2616 | 3.10E-01 | **2.38E-06** |
| **PANp** | 0.5941 | 4.61E-01 | -1.5808 | 7.34E-02 | -2.175 | **1.18E-02** | **1.55E-02** |
| **PANn** | 6.4249 | **2.72E-11** | 1.8623 | 6.98E-02 | -4.5625 | **1.52E-05** | **1.82E-10** |
| **PANg** | 3.3873 | **8.30E-04** | -2.2114 | 4.51E-02 | -5.5987 | **2.59E-08** | **1.47E-06** |
| **PANs** | -0.2023 | 4.07E-01 | -0.4986 | 6.23E-02 | -0.2964 | 2.10E-01 | 2.47E-01 |
| **PANtotal** | 10.204 | **2.58E-05** | -2.429 | 3.55E-01 | -12.633 | **2.69E-07** | **1.93E-07** |
| **Course** | 1.3135 | **2.96E-07** | 0.608 | **2.63E-02** | -0.7545 | **5.75E-03** | **4.47E-06** |

**Table 3.** Enrichment of gene-set (which identified by cluster analysis) for psychiatric GWAS results

| Clusters | SCZ | Bipolar | Depression |
| --- | --- | --- | --- |
| Cluster One | 0.087 | 0.496 | 0.399 |
| Cluster Two | 0.100 | **0.013** | 0.207 |
| Cluster Three | **0.017** | 0.174 | 0.405 |

**Fig. 4.**
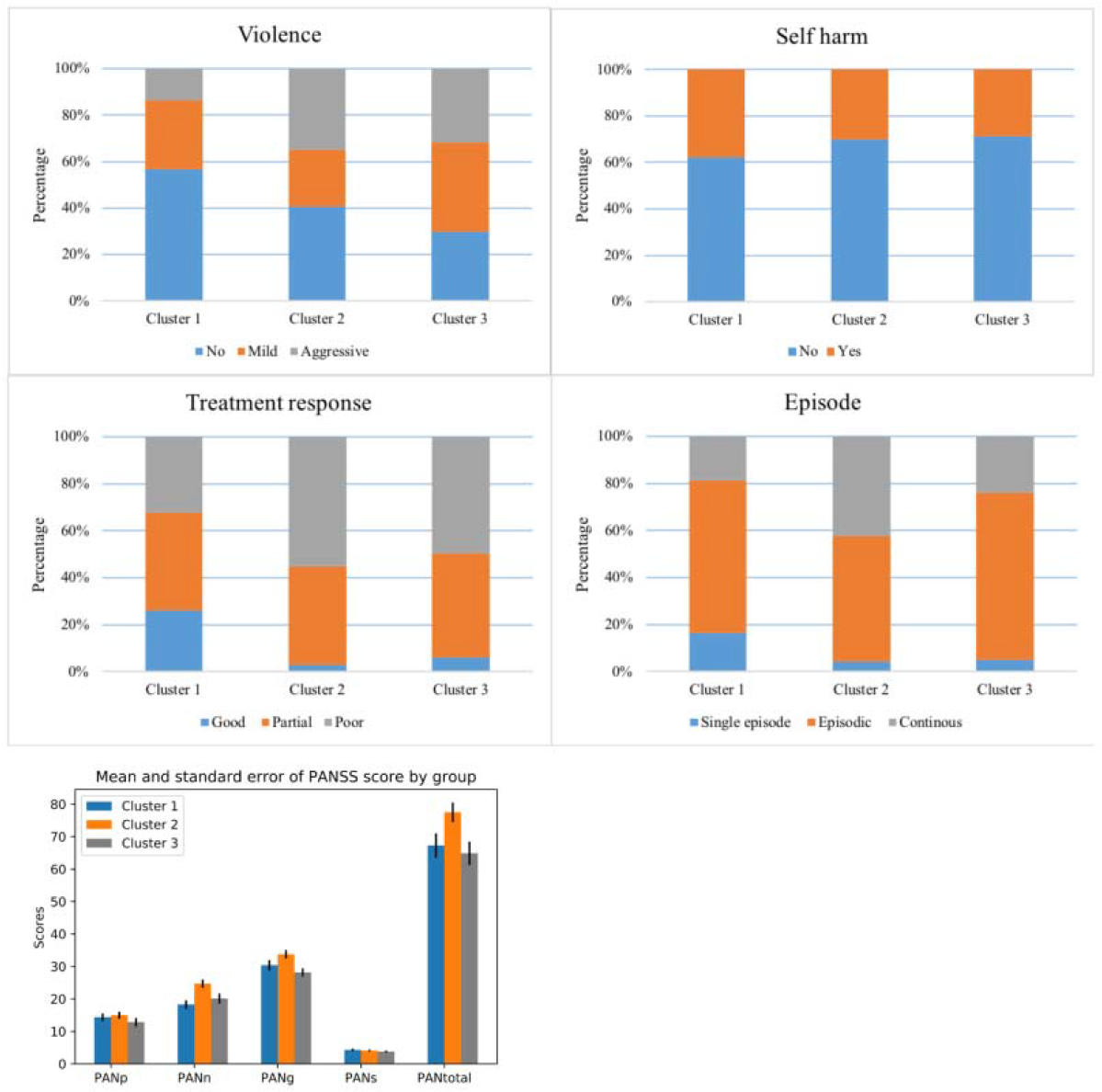
Comparison across outcome-related variables by subgroups for SCZ patients

Notably, our approach significantly outperformed clustering based on random assignment (p<0.001, 1000 Monte-Carlo simulations). To further assess clusters are genuinely present in the dataset, we also applied “sigclust” to our SCZ data which is also statistically significant (p= 0 as reported by sigclust).

#### Gender stratified analysis

We observed significant difference on gender ratio in the 3 identified subgroups (Fig. 3). In this regard, we conducted further analysis to examine whether the observed associations with clinical outcomes were purely driven by gender differences.

Firstly, we excluded all females in cluster 1, and repeated the association analyses on input and outcome variables considering male subjects only. As expected, we still observed significant differences across most outcome features, except for violence, self-harm and PANSS aggression subscale score (Table S3, Supplementary Fig. 1 and Fig. 2).

Subsequently, we repeated our multi-view clustering analysis method on *female* patients only. The best solution consisted of 3 subgroups. Similar with male-only analysis, we again observed significant differences across most clinical outcomes including treatment response, 3 PANSS subscale scores (negative, general and total score) as well as disease course (Table S4, Supplementary Fig. 3, and Fig. 4).

#### Selected genes and pathway analysis

Among selected genes by clustering analysis, numerous were involved in schizophrenia or related pathophysiological processes, such as *ZNF804A*^36,37^, *SNX19*^38^, *LRP1*^39^, *CACNB*2^40,41,42^. *ZNF804A* has been identified as a top risk gene in schizophrenia which is implicated in neurodevelopmental processes^43^. In addition, we examined whether the genes selected by the cluster algorithm were ‘enriched’ for GWAS hits. We tested whether the selected genes as a whole had lower *p-*values from GWAS (of SCZ, bipolar disorder and depression) than those not selected. As expected, the strongest enrichment was observed for SCZ, and significant enrichment was also observed for bipolar disorder (cluster 2). Note that the biclustering algorithm selected these genes ‘blindly’, as no pre-screening for association with SCZ was performed. We also note that the genes that characterized SCZ prognosis and clinical features may *not* be the same as those that affect susceptibility to the disease, but we expect a partial overlap.

As we employed a gene-based approach in our analysis, we may characterize each subgroup by involved genes and pathways readily. The top enriched biological pathways and gene ontology are demonstrated in Table S7; we highlight a few pathways here. For cluster 1, antigen processing and presentation^44^, generation of second messenger molecules^45,46^, autoimmune thyroid disease^47^, pyrimidine metabolism^48^ were among the top pathways. For the second cluster, some of the involved pathways included arachidonic acid metabolism^49,50^, glutathione conjugation^51,52^, glutathione-mediated detoxification^53^ and others. As for the third cluster, some significant pathways included DNA damage reversal^54,55^, Vitamin D3 (cholecalciferol) metabolism^56^, metabotropic glutamate/pheromone receptors^57,58^. Some enriched pathways were shared among different clusters while some were not. Notably, numerous enriched pathways were associated with psychiatric disorders or brain functioning. Please refer to the attached references for potential relationships of the named pathways with SCZ or other psychiatric disorders. For example, immune and inflammatory processes are postulated to play an important role in SCZ pathogenesis^59^ and ‘antigen processing and presentation’ was the second most significantly enriched pathway (*p*=1.08E-10) in a SCZ GWAS^44^; increased breakdown of arachidonic acid was revealed to be responsible for neuronal deficits in SCZ^49^; DNA damage and dysfunctional DNA repair^60^ has been reported to contribute to the pathophysiology of psychiatric disorders^54^; glutamatergic dysfunction has been implicated in SCZ and proposed as targets for new drugs^61^. For the tissue specificity analysis, we observed *all* sets of selected genes in all tissues were significantly enriched in DEGs in the *brain* (Supplementary Fig. 5).

**Fig. 5.**
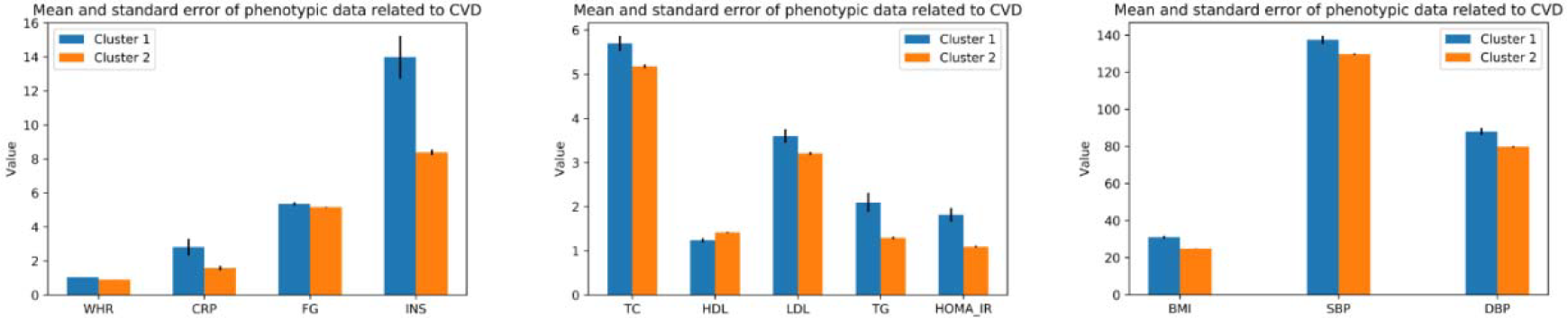
Comparison across input clinical features by male subgroups from multi-view clustering

### Application to Northern Finland Birth Cohort

Next we applied the proposed framework to the NFBC cohort to stratify patients into different levels of cardiometabolic risks. There exists significant gender differences in terms of risk factors, prevalence, age at onset and clinical presentation of CVD^62,63^. The Framingham scoring system and criteria for metabolic syndrome are also set separately for males and females ^64^. In view of the well-established differences between genders, we performed the analysis separately in males and females.

#### Gender stratified analysis

Firstly, we applied our framework to male subjects only. The best solution consisted of two subgroups, with 146 and 2306 subjects in each subgroup. We observed highly significant differences among all 12 input clinical variables (Fig.5, Table 4) between two subgroups. The best solution comprised two clusters (‘high CV risk’ and ‘low CV risk’) with marked differences in cardiometabolic risk factors. More specifically, subjects in the first subgroup had higher levels of LDL, TG, TC, BP, CRP, fasting glucose/insulin and were more obese (average BMI of 31.03). Subjects in the second subgroup showed a more favourable profile of cardiovascular risks.

**Table 4.** Differences in input clinical variables among subgroups derived from cluster analysis of only ***males***

| Measures | Estimate | P value |
| --- | --- | --- |
| **WHR** | -0.136587 | 1.32E-199 |
| **CRP** | -1.2323 | 7.34E-06 |
| **FG** | -0.19770 | 1.27E-04 |
| **INS** | -5.5978 | 1.72E-51 |
| **TC** | -0.51332 | 1.92E-09 |
| **HDL** | 0.17767 | 1.43E-10 |
| **LDL** | -0.39540 | 2.13E-07 |
| **TG** | -0.8068 | 2.97E-31 |
| **HOMA-IR** | -0.72122 | 1.27E-52 |
| **BMI** | -6.1981 | 1.77E-100 |
| **SBP** | -7.762 | 3.52E-13 |
| **DBP** | -8.2871 | 6.93E-18 |

WHR, waist-hip ratio; CRP, C-reactive protein; FG, fasting glucose; INS, fasting insulin; TC, total cholesterol; HDL, high-density cholesterol; LDL, low-density cholesterol; TG, triglyceride; HOMA-IR, Homeostatic Model Assessment for Insulin Resistance; BMI, body mass index; SBP, systolic blood pressure; DBP, diastolic blood pressure.

We also computed prediction strength (*ps*) for our male-only clustering results, and obtained a *ps* of 0.759 for our selected solution, signifying relatively good ability for the clustering results to be generalized to a new dataset. To further verify the reliability of our solution, we compared the *ps* with that obtained from a random clustering approach. The solution is significantly better than by chance (p<0.001, 1000 random cluster assignments). “sigclust” also returned p-value of 0, indicating existence of genuine cluster structure in the data instead of random sampling variations.

We repeated our approach on *female* subjects of NFBC. The best solution was composed of 2 subgroups with 65 and 2465 subjects respectively. Again, we observed significant differences among all 12 input clinical variables (Fig. 6, Table 5). Compared with subjects in the 2^nd^ subgroup, those in the 1^st^ subgroup manifested significantly higher levels of cardiometabolic risk factors in all input clinical variables except HDL.

**Table 5.** Differences in input clinical variables among subgroups derived from cluster analysis of only ***females***

| Measures | Estimate | P value |
| --- | --- | --- |
| **WHR** | -0.284545 | 3.87E-224 |
| **CRP** | -2.4734 | 1.45E-06 |
| **FG** | -0.19036 | 1.80E-03 |
| **INS** | -5.7274 | 4.17E-34 |
| **TC** | -0.3737 | 9.20E-04 |
| **HDL** | 0.26044 | 1.60E-08 |
| **LDL** | -0.41459 | 2.91E-05 |
| **TG** | -0.43240 | 1.26E-11 |
| **HOMA-IR** | -0.72882 | 2.87E-33 |
| **BMI** | -9.3350 | 5.28E-59 |
| **SBP** | -10.482 | 8.05E-12 |
| **DBP** | -8.076 | 9.36E-10 |

**Fig. 6.**
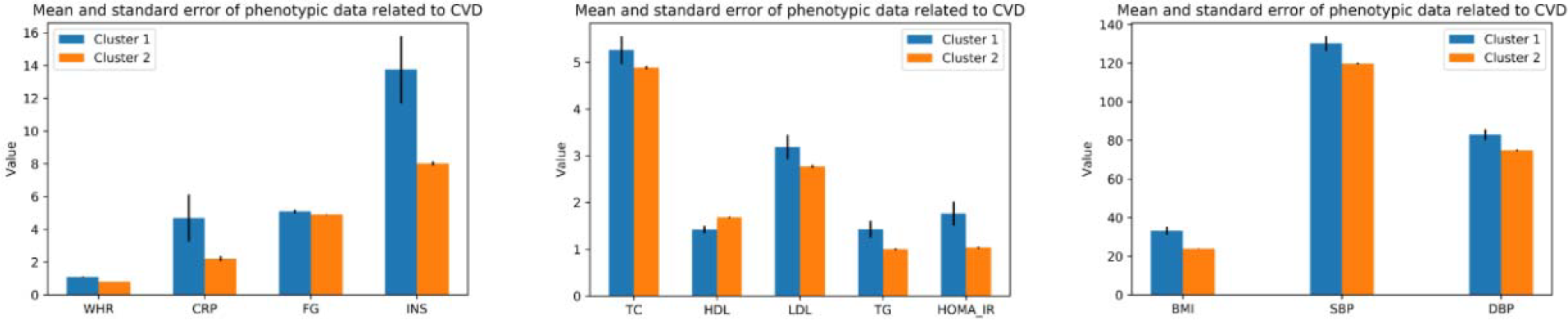
Comparison across input clinical features by female subgroups from multi-view clustering

We also employed prediction strength to evaluate the validity of our approach. For our selected solution, we got a *ps* of 0.826, indicating good clustering performance. Our approach significantly outperformed randomly assigned clustering method (p<0.001, 1000 random cluster assignments). Also, we applied “sigclust” to our female-only dataset, which confirmed the presence of cluster structure in our data (p=0).

#### Selected genes and pathway analysis

We separately analysed the selected genes from male-only and female-only clustering analysis. Numerous selected genes were implicated in CVD or related pathophysiological processes, including *CEP68*^65^, *SREBF*^66^, *FMO*^67^, *ITGB*^68^ etc. For example, *CEP68* has been identified as a top risk gene for elevated blood pressure. We also examined whether these selected genes are enriched for GWAS ‘hits’ of cardiometabolic disorders. In brief, we first performed gene-based test on CAD and DM GWAS data, then examined whether the selected genes from cluster analysis have lower *p*-values than the non-selected genes. We observed this is indeed true for both female-only and male-only clustering analysis (Table 6). The results provide support for the validity of our approach, as the cluster algorithm is ‘blind’ to which genes being associated with CAD or DM beforehand.

**Table 6.**
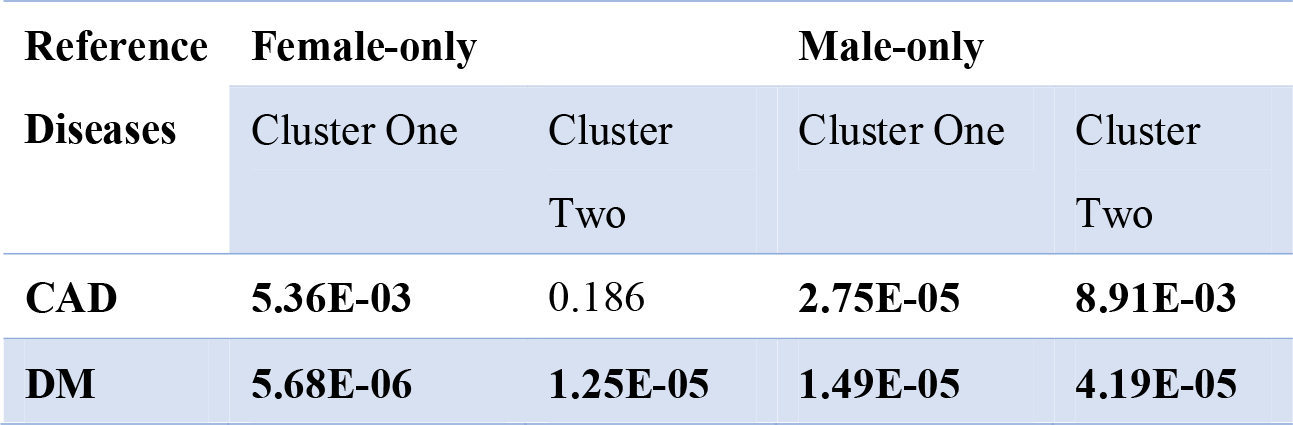
Enrichment of genes identified by the cluster analysis for CAD (coronary artery disease) and DM (diabetes mellitus) GWAS results

| Reference<br>Diseases | Female-only |  | Male-only |  |
| --- | --- | --- | --- | --- |
|  | Cluster One | Cluster Two | Cluster One | Cluster Two |
| **CAD** | **5.36E-03** | 0.186 | **2.75E-05** | **8.91E-03** |
| **DM** | **5.68E-06** | **1.25E-05** | **1.49E-05** | **4.19E-05** |

Also, we analysed the top enriched biological pathways for males and females respectively (as demonstrated in Table S8 and Table S9). To highlight a few potentially interesting pathways, for female-only subjects, the involved pathways of cluster 1 included NRF2 pathway^69^, Pathways in Pathogenesis of Cardiovascular Disease and Proteasome Degradation^70^; for cluster 2, Arrhythmogenic Right Ventricular Cardiomyopathy^71^, NRF2 pathway and Adipogenesis^72^ were among the top enriched pathways. As for male-only subjects, the top enriched pathways for cluster 1 included Tamoxifen metabolism^73^, Complement Activation^74^ and BDNF-TrkB Signaling^75^; the involved pathways for the second cluster included Apoptosis Modulation and Signaling^76^, Cardiac Hypertrophic Response^77^ and NRF2 pathway. As expected, some of the top involved pathways were shared among female-only and male-only clusters while others were not. Notably, some of the top enriched pathways were associated with increased risk of cardiovascular disorders, for example, NRF2 pathway plays a significant role in the development and progression of CVD^69^. Apoptosis is shown to be involved in the development of both acute and chronic heart failure^76^. For details about the top enriched pathways and gene ontology, please refer to Table S8 and S9. Finally, for the tissue specificity analysis, significant enrichment in heart, liver and artery were observed in both females and males (Supplementary Fig. 6 and Fig. 7).

## Discussion

In this study, we have presented a novel framework capable of discovering latent subgroups of complex disease by leveraging patients’ clinical and GWAS-predicted expression profiles. We verified the feasibility and validity of our proposed approach by applying it to two different datasets. For example, in the SCZ dataset, the derived subgroups showed significant differences in disease outcomes such as treatment response, course of illness and symptom scores. In addition, we observed satisfactory prediction strength (the ability of the clustering model to ‘predict’ clusters in a new dataset) for both applications in SCZ and cardiometabolic disorders. Moreover, we found that the genes ‘blindly’ selected by the cluster algorithm are significantly enriched for those discovered in genetic association studies of SCZ and cardiovascular diseases, supporting the biological relevance of the clustering approach.

To our knowledge, this is the first study to leverage GWAS-predicted expression profiles and clinical variables to discover complex disease subgroups. Through imputation to expression levels, GWAS data might be readily analysed using other forms of clustering techniques, such as those developed for subtyping oncology patients. Therefore, our proposed analytic framework is highly extensible to current or even future unsupervised learning or clustering methodologies. In addition, our proposed approach can be applied to any existing GWAS datasets, which are often of much larger sample sizes compared to expression studies. As we have mentioned in the introduction, there are numerous other advantages of the presented framework. Since genetic variations have been mapped to expression levels, the discovered subgroups are likely more biologically relevant and interpretable than a pure SNP-based analysis. Another important advantage is that we can easily extend to multiple tissues, especially those that are difficult to access (e.g. brain). The analysis results are also unlikely to be confounded by other factors such as medication use. As such, differences among derived subgroups will not be merely due to differences in the drugs prescribed or intervention given. Imputation of SNP data to gene-level data also reduces the dimension substantially, increasing the computational speed and ease of analysis. For example, for the SCZ data, the computation efficiency is dramatically improved by employing a gene-based compared to a SNP-based approach to clustering, while the gene-based approach still being able to divide the patients into diverse subgroups with significantly different prognosis (Table 7).

**Table 7.**
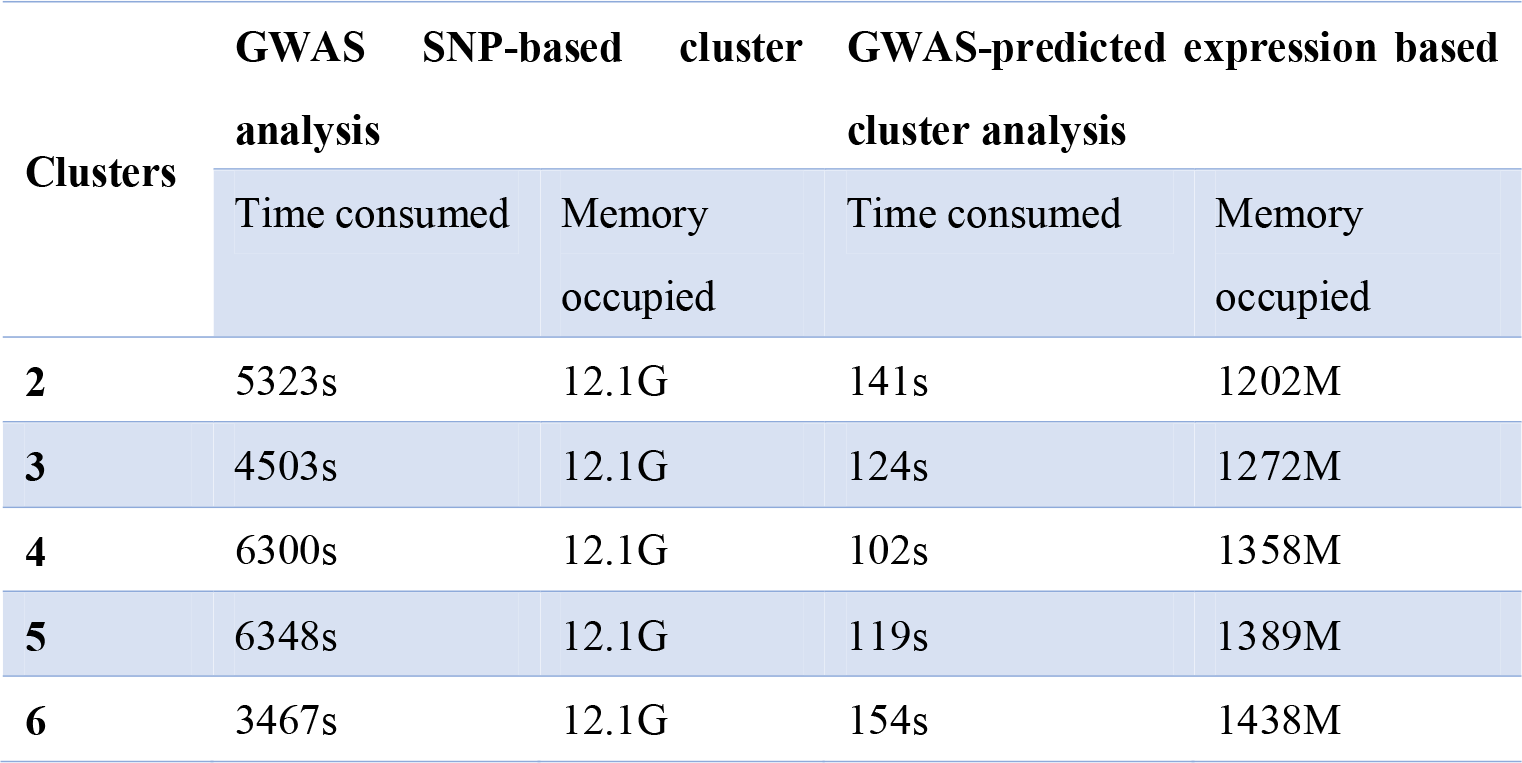
Comparison on computational cost across different methods

The purpose of our study is to find clinically and biologically homogenous subgroups of patients. However this similarity may extend beyond the outcome variables collected in our dataset, for example predisposition to comorbidities or complications, response or side-effects to current or even new medications etc. The clinical implications of the derived clusters may therefore be beyond the variables recorded. In this regard, one limitation of the current study is that some of the outcome variables are not available. For instance, for the NFBC 1966 dataset, we do not have longitudinal data on the cardiovascular outcomes (e.g. CAD, stroke, CVD deaths); such information will be valuable in testing whether derived subgroups of subjects differed in cardiovascular outcomes in the long run. In a similar vein, the clinical data used as *input* for clustering are also not complete. For example, the neurocognitive profiles collected for SCZ patients are not thorough and apart from CRP the NFBC dataset do not have other measurement of serum biomarkers. Another limitation is that expression imputation is based on the GTEx dataset, which is of modest sample size. The imputation is subject to error and some genes may not be predicted as accurately as others. The imputation may be less than optimal for non-Caucasian populations, due to nature of the GTEx dataset in which ~85% are Caucasians. Nevertheless, empirically we observed reasonable performance of our clustering framework, and we believe the situation might improve when larger genotype-transcriptome studies are released in the future. A related open question is how to accommodate imputed expression from different tissues. One solution, as we employed here, is to extract the most relevant tissues and model these tissues only. Methods for prioritizing the most important tissues for a disease are emerging^28^. However, it remains unknown whether this is the most optimal approach. For example, it may be possible to model more tissues but to assign a weighting according to the relevance of each tissue to the disorder.

One concern of cluster analysis using genomic data is the effect of population stratification. Population stratification is a confounder in genetic association studies, for which the aim is to uncover susceptibility variants for a trait. However, in a cluster analysis context, we argue that population stratification is usually *not* a major problem, particularly from a clinical point of view. We have discussed this issue in detail in our previous work^8^. Briefly, from a clinical perspective, the clustering is satisfactory if patients can be divided into groups of *clinical differences*, for example different prognosis, survival or drug responses. If patients are clustered into different groups due to or partially due to (possibly subtle) ancestry differences, as long as the subgroups are clinically diverse, this clustering is still useful and valid from a clinical viewpoint. There are two possibilities for diverse clinical profiles in different ethnic groups. The ethnic difference may be associated with other environmental factors (e.g. socioeconomic background, dietary/lifestyle patterns) that are also linked to the disease profiles or prognosis. In this case, population stratification can be “beneficial” as the clustering framework can consider extra information captured by the ethnic differences. The model can be considered ‘valid’ as long as it is applied to a similar population. However, if we only wish to reveal the genes contributing to the disease subgroups, the genetic variants identified may not have direct biological relevance to the studied disease under this condition. In this study, the SCZ dataset is exclusively collected in Hong Kong while the NFBC sample is from Finland only. We observed significant enrichment of the selected genes for susceptibility genes of SCZ/CAD/DM in other GWAS, and also revealed pathways of functional importance, indicating the selected genes may indeed be biologically relevant, although further functional studies are required to confirm the findings.

Another possibility (which could co-exist with the first), is that some variants that are different among the ethnic groups are also *biologically* related with the disease. For example, an ethnic subgroup may have a higher/lower frequency of certain variant(s) affecting drug metabolism leading to better/worse response. The clustering is clearly valid in this scenario.

For the NFBC example, we observed imbalance in the derived subgroups in which the ‘high-risk’ subgroup contains a small number of subjects only. This is probably reasonable as all subjects are relatively young (aged 31) and the proportion of subjects having high CV risks is likely to be low. Interestingly, we computed that the proportion of NFBC participants with clinically defined metabolic syndrome (according to the latest Harmonized criteria^31^) is 5.5% in males and 2.5% in females. These numbers are close to the proportion of subjects in the ‘high risk’ cluster using our proposed clustering framework. This suggests the ‘imbalance’ in the derived clusters makes clinical sense, despite we do not give the algorithm a priori guidance on the distribution of the clusters. One concern may be that whether the subgroups derived from the present clustering framework will be very similar to those derived by existing criteria of MetS, given the similar proportion of subjects subtyped as ‘high risk’. If this is the case, there is not much added value of including genomic data. We checked that the derived cluster from our approach have only *partial* overlap (21.1% for males, 16.9% for females) with the existing criteria for MetS, suggesting that genomic data adds to existing clinical information and provides an alternative, biologically-driven approach to characterize patient subgroups with high cardiometabolic risks. Intuitively, genetic data reflects the predisposition to develop certain traits or diseases, and may help predict the *future* risk of MetS or CVD, as opposed to existing approaches which only rely on cardiometabolic parameters measured at present. For instance, a young subject may not have MetS yet but maybe genetically predisposed to developing MetS and CVD; these subjects may be picked up by the proposed subtyping approach via integrating genomic and clinical data.

In oncology, studies on cancer subtyping have greatly benefited from resources of genomic data such as TCGA. Some approaches to cancer subtyping have also observed clinical applications ^78^. It is worth noting that many of these studies and methodologies developed on cancer subtyping utilized expression data. From a broader perspective, our presented approach which leverages transcriptome data are linked to these works, as clustering methodologies developed in cancer research (that utilize expression profiles) could be ‘translated’ to other complex diseases under our presented framework.

To summarize, we proposed a novel analytic framework to uncover subtypes of complex diseases by leveraging both clinical and GWAS-imputed expression profiles. The derived subgroups exhibited significant differences across numerous outcome variables and/or showed good prediction strength, indicating the feasibility and validity of our proposed method. Enrichment of genes selected by the cluster algorithm for GWAS hits provided further support to our approach. From a clinical point of view, stratification of patients is crucial in provided more targeted prevention as well as intervention strategies; from a more basic science perspective, our approach may help identify subtype-specific biological pathways and processes, and the development of more personalized drug therapies for patients.

## Acknowledgements

This work was supported partially by the Lo Kwee Seong Biomedical Research Fund, a Direct Grant from The Chinese University of Hong Kong, and RGC Collaborative Research Fund C4054-17WF. We are grateful to Dr. Eric Cheung, Dr. Emily Wong, Dr. Ronald Chen, Prof. Tao Li, Prof. Eric Chen, Tomy Hui for their help in subject recruitment and Prof. Miaoxin for help in GWAS analysis in the schizophrenia dataset. We also thank Prof. Stephen Tsui for computing support.

