## Supplementary material for "Uncovering complex disease subtypes by integrating clinical data and imputed transcriptome from genome-wide association studies: Applications in psychiatry and cardiovascular medicine": Supp Figures

**Supplementary Figures**

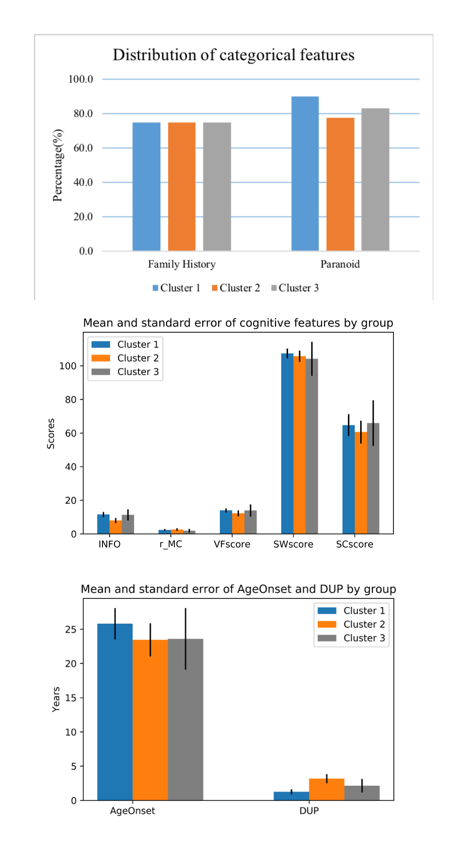

Supplementary Fig. 1 Comparison of input clinical features across *female* SCZ patient subgroups

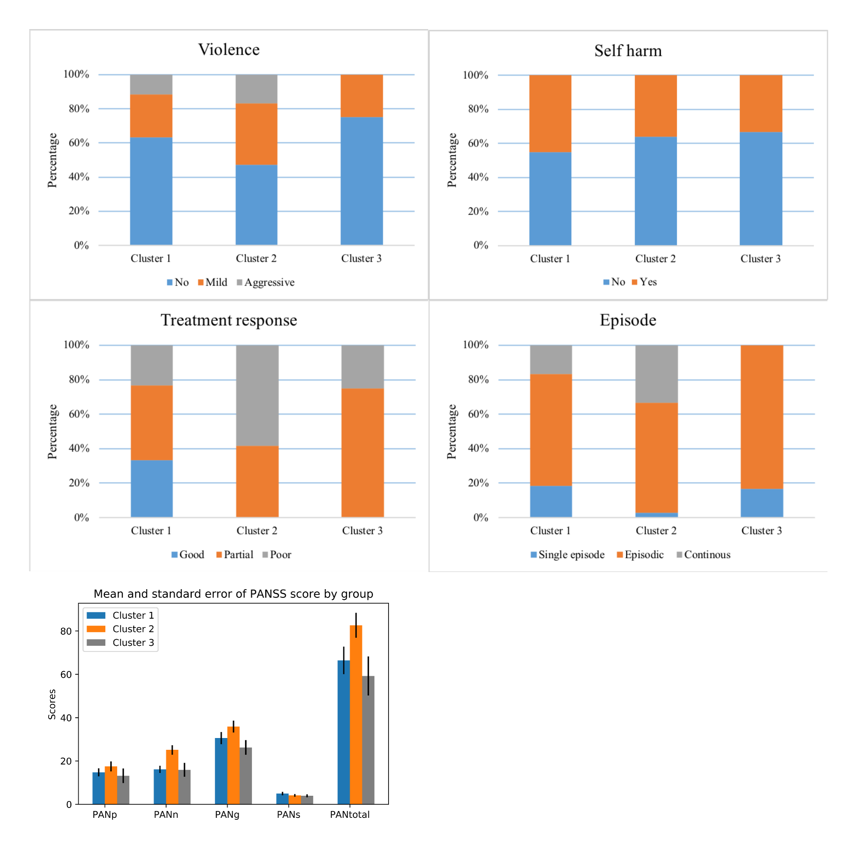

Supplementary Fig. 2 Comparison of outcome variables across *female* SCZ patient subgroups

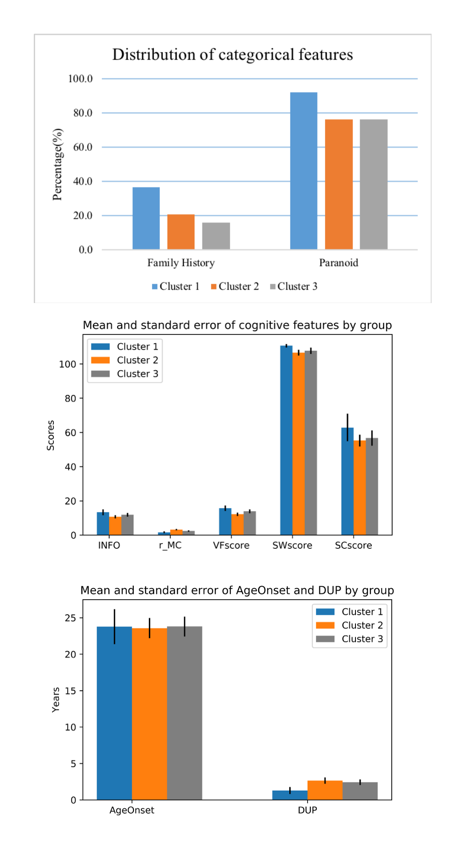

Supplementary Fig. 3 Comparison of input clinical features across *male* SCZ patient subgroups

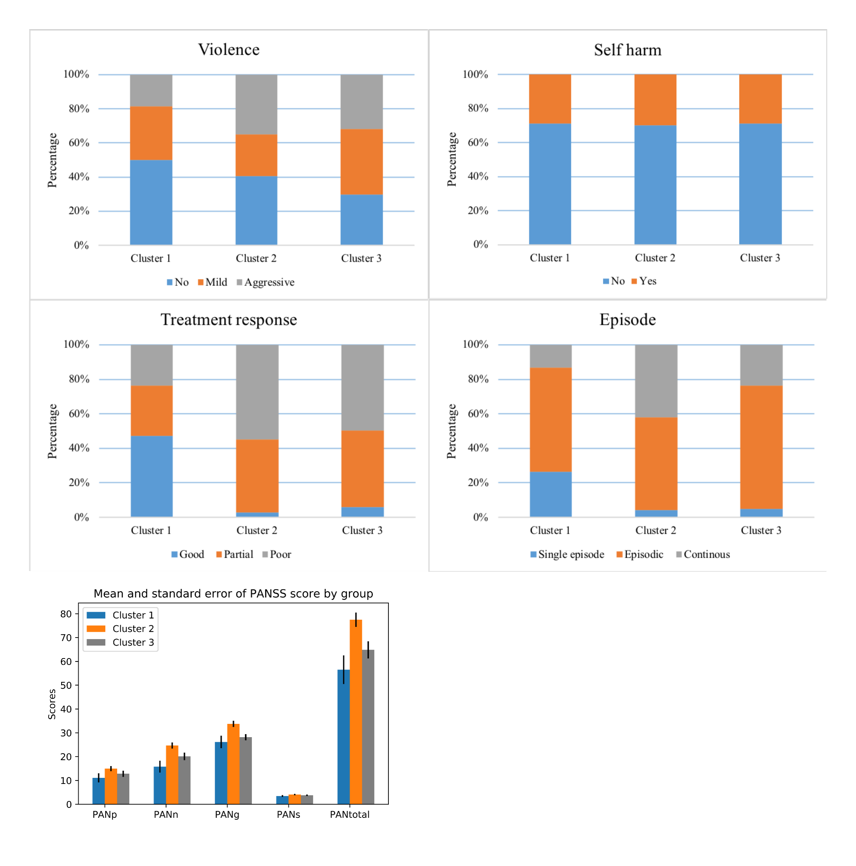

Supplementary Fig. 4 Comparison of outcome variables across *male* SCZ patient subgroups

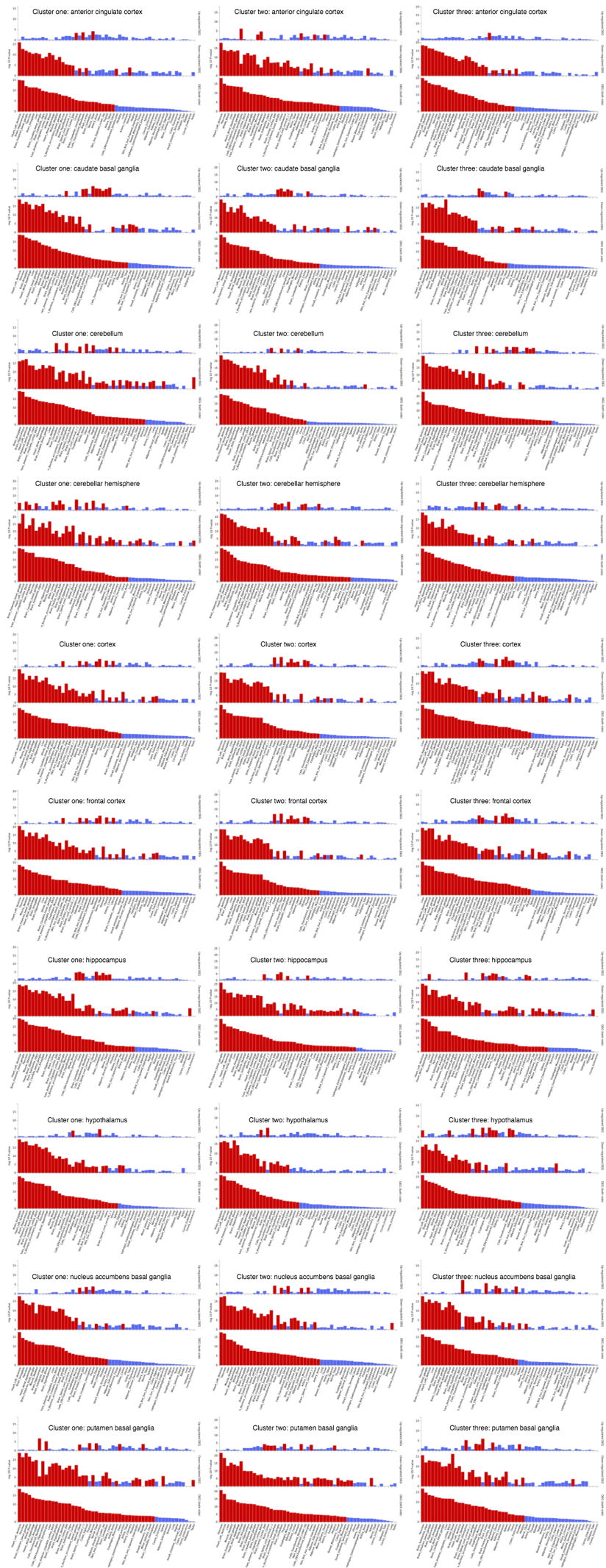

Supplementary Fig. 5 enrichment for DEGs across 53 specific tissues for selected genes by groups of SCZ patients

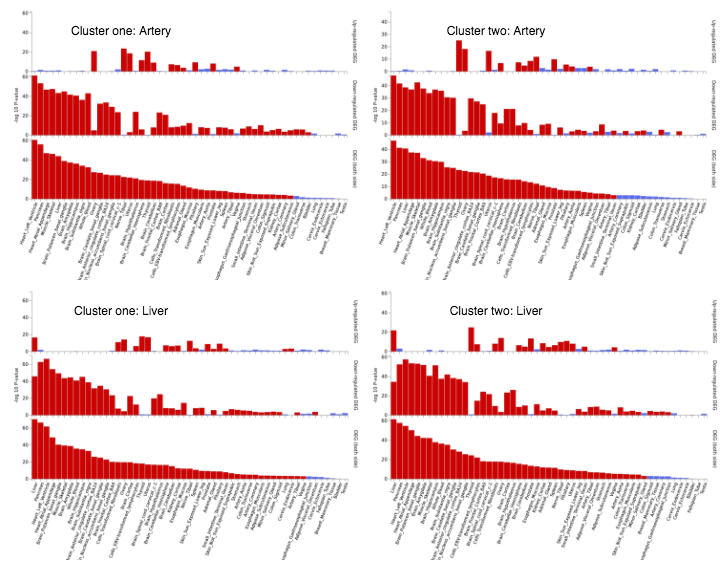

Supplementary Fig. 6 enrichment for DEGs across 53 specific tissues for selected genes by female groups of CVD subjects

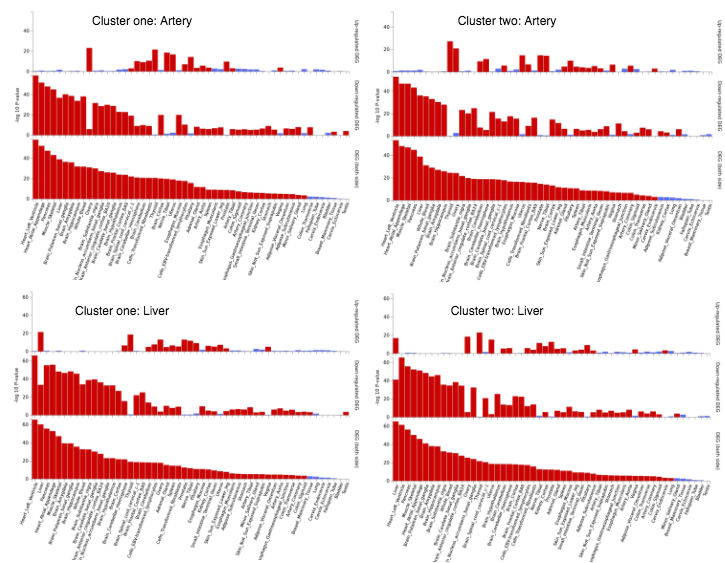

Supplementary Fig. 7 enrichment for DEGs across 53 specific tissues for selected genes by male groups of CVD subjects
