## Supplementary material for "Uncovering complex disease subtypes by integrating clinical data and imputed transcriptome from genome-wide association studies: Applications in psychiatry and cardiovascular medicine": Supp tables

**Supplementary Tables**

Supplementary Table S1 Comparison of input features across schizophrenia (SCZ) patient subgroups (as identified by the biclustering algorithm)

| Features | Cluster1 VS 2 | | | Cluster 1 VS 3 | | | Cluster 2 VS 3 | | | Overall | |
| --- | --- | --- | --- | --- | --- | --- | --- | --- | --- | --- | --- |
|  | Estimate | P values | Q values | Estimate | P values | Q values | Estimate | P values | Q values | P values | Q values |
| FHxMI | -0.4018 | 0.1486 | 2.67E-01 | -0.7297 | 2.65E-02 | *6.81E-02* | -0.3278 | 3.39E-01 | 5.09E-01 | 6.39E-02 | *1.28E-01* |
| Dx_type | 0.7234 | 2.23E-02 | *6.69E-02* | 0.734 | 3.05E-02 | *6.86E-02* | 0.0969 | 1.06E-02 | **3.82E-02** | 3.29E-02 | *6.97E-02* |
| INFO | -0.3882 | 5.71E-01 | 6.81E-01 | 0.7716 | 3.04E-01 | 4.99E-01 | 1.1598 | 1.21E-01 | 2.29E-01 | 3.06E-01 | 4.99E-01 |
| r_MC | 1.0219 | 1.53E-04 | **5.51E-03** | 0.1784 | 5.42E-01 | 6.73E-01 | -0.8436 | 6.38E-03 | **3.28E-02** | 3.68E-04 | **6.62E-03** |
| VFscore | -1.85714 | 5.28E-03 | **3.17E-02** | -0.07921 | 9.13E-01 | 9.13E-01 | 1.7779 | 2.02E-02 | *6.61E-02* | 9.11E-03 | **3.64E-02** |
| SWscore | -0.8845 | 4.63E-01 | 6.31E-01 | 0.2053 | 8.76E-01 | 9.01E-01 | 1.0898 | 4.03E-01 | 5.80E-01 | 6.60E-01 | 7.20E-01 |
| SCscore | -8.052 | 2.99E-03 | **2.69E-02** | -6.575 | 2.64E-02 | *6.81E-02* | 1.477 | 6.06E-01 | 6.82E-01 | 7.46E-03 | **3.36E-02** |
| AgeOnset | -0.9388 | 3.19E-01 | 4.99E-01 | -0.7088 | 4.91E-01 | 6.31E-01 | 0.23 | 8.22E-01 | 8.70E-01 | 5.86E-01 | 6.81E-01 |
| DUP | 0.8312 | 1.22E-03 | **1.46E-02** | 0.6131 | 2.87E-02 | *6.86E-02* | -0.2181 | 4.76E-01 | 6.31E-01 | 3.82E-03 | **2.75E-02** |

Supplementary Table S2 Comparison of outcome-related variables across SCZ patient subgroups

| Features | Cluster1 VS 2 | | | Cluster 1 VS 3 | | | Cluster 2 VS 3 | | | Overall | |
| --- | --- | --- | --- | --- | --- | --- | --- | --- | --- | --- | --- |
|  | Estimate | P values | Q values | Estimate | P values | Q values | Estimate | P values | Q values | P values | Q values |
| Violence | 0.8576 | 1.70E-04 | **3.83E-04** | 1.03601 | 1.88E-05 | **5.21E-05** | -0.1731 | 4.68E-01 | 4.81E-01 | 1.09E-05 | **3.57E-05** |
| Self-harm | -0.3438 | 1.71E-01 | 2.26E-01 | -0.4058 | 1.45E-01 | 2.01E-01 | -0.0621 | 8.29E-01 | 8.29E-01 | 2.45E-01 | 2.94E-01 |
| Treatment response | 1.2489 | 1.27E-07 | **9.14E-07** | 1.0106 | 6.94E-05 | **1.67E-04** | -0.2616 | 3.10E-01 | 3.60E-01 | 1.20E-07 | **9.14E-07** |
| PANp | 0.5941 | 4.61E-01 | 4.81E-01 | -1.5808 | 7.34E-02 | ***1.06E-01*** | -2.175 | 1.18E-02 | **2.24E-02** | 4.63E-02 | **7.58E-02** |
| PANn | 6.4249 | 2.72E-11 | **9.79E-10** | 1.8623 | 6.98E-02 | ***1.05E-01*** | -4.5625 | 1.52E-05 | **4.56E-05** | 9.42E-11 | **1.70E-09** |
| PANg | 3.3873 | 8.30E-04 | **1.76E-03** | -2.2114 | 4.51E-02 | ***7.58E-02*** | -5.5987 | 2.59E-08 | **3.11E-07** | 2.34E-06 | **8.42E-06** |
| PANs | -0.2023 | 4.07E-01 | 4.44E-01 | -0.4986 | 6.23E-02 | ***9.75E-02*** | -0.2964 | 2.10E-01 | 2.61E-01 | 1.76E-01 | 2.26E-01 |
| PANtotal | 10.204 | 2.58E-05 | **6.63E-05** | -2.429 | 3.55E-01 | 3.99E-01 | -12.633 | 2.69E-07 | **1.52E-06** | 1.25E-06 | **5.00E-06** |
| Course | 1.3135 | 2.96E-07 | **1.52E-06** | 0.608 | 2.63E-02 | **4.73E-02** | -0.7545 | 5.75E-03 | **1.15E-02** | 8.74E-07 | **3.93E-06** |

Supplementary Table S3 Comparison of outcome-related features across SCZ *male* patient subgroups

| Features | Cluster1 VS 2 | | | Cluster 1 VS 3 | | | Cluster 2 VS 3 | | | Overall | |
| --- | --- | --- | --- | --- | --- | --- | --- | --- | --- | --- | --- |
|  | Estimate | P values | Q values | Estimate | P values | Q values | Estimate | P values | Q values | P values | Q values |
| Violence | 0.565 | 9.99E-02 | 1.50E-01 | 0.7396 | 3.60E-02 | 5.89E-02 | -0.1731 | 4.68E-01 | 5.27E-01 | 1.06E-01 | 1.51E-01 |
| Self-harm | 0.05064 | 9.00E-01 | 9.53E-01 | -0.01143 | 9.78E-01 | 9.78E-01 | -0.0621 | 8.29E-01 | 9.04E-01 | 9.75E-01 | 9.78E-01 |
| Treatment response | 2.3499 | 8.81E-09 | **5.29E-08** | 2.1024 | 4.69E-07 | **1.53E-06** | -0.2616 | 3.10E-01 | 3.65E-01 | 1.41E-08 | **7.25E-08** |
| PANp | 3.852 | 1.33E-03 | **3.19E-03** | 1.677 | 1.76E-01 | 2.26E-01 | -2.175 | 1.18E-02 | **2.12E-02** | 1.51E-03 | **3.40E-03** |
| PANn | 8.882 | 2.85E-09 | **2.05E-08** | 4.319 | 4.41E-03 | **8.82E-03** | -4.5625 | 1.52E-05 | **3.91E-05** | 8.71E-10 | **1.05E-08** |
| PANg | 7.599 | 8.39E-08 | **3.36E-07** | 2.001 | 1.65E-01 | 2.20E-01 | -5.5987 | 2.59E-08 | **1.17E-07** | 2.24E-10 | **4.03E-09** |
| PANs | 0.6312 | 4.88E-02 | 7.64E-02 | 0.3348 | 3.14E-01 | 3.65E-01 | -0.2964 | 2.10E-01 | 2.61E-01 | 1.09E-01 | 1.51E-01 |
| PANtotal | 20.964 | 1.64E-09 | **1.48E-08** | 8.332 | 1.78E-02 | **3.05E-02** | -12.633 | 2.69E-07 | **9.68E-07** | 5.23E-11 | **1.88E-09** |
| Course | 2.0196 | 2.51E-06 | **6.95E-06** | 1.2908 | 2.77E-03 | **5.87E-03** | -0.7545 | 5.75E-03 | **1.09E-02** | 2.22E-06 | **6.66E-06** |

Supplementary Table S4 Comparison of outcome-related features across SCZ *female* patient subgroups

| Features | Cluster1 VS 2 | | | Cluster 1 VS 3 | | | Cluster 2 VS 3 | | | Overall | |
| --- | --- | --- | --- | --- | --- | --- | --- | --- | --- | --- | --- |
|  | Estimate | P values | Q values | Estimate | P values | Q values | Estimate | P values | Q values | P values | Q values |
| Violence | 0.6918 | 9.44E-02 | 1.79E-01 | 0.2547 | 6.84E-01 | 7.24E-01 | 0.3823 | 4.47E-01 | 5.19E-01 | 2.46E-01 | 3.41E-01 |
| Self-harm | -0.2513 | 5.58E-01 | 6.20E-01 | -0.8979 | 2.09E-01 | 3.14E-01 | -0.8293 | 9.31E-02 | 1.79E-01 | 4.05E-01 | 4.92E-01 |
| Treatment response | 1.4548 | 5.75E-04 | **2.96E-03** | 0.7333 | 1.99E-01 | 3.14E-01 | 2.1656 | 3.20E-05 | **1.04E-03** | 1.78E-03 | **7.12E-03** |
| PANp | 2.4167 | 1.10E-01 | 1.98E-01 | -2.6667 | 2.39E-01 | 3.41E-01 | 6.662 | 9.57E-05 | **1.04E-03** | 7.61E-02 | 1.61E-01 |
| PANn | 6.1611 | 1.15E-04 | **1.04E-03** | 0.2167 | 9.25E-01 | 9.25E-01 | 5.931 | 1.44E-03 | **6.48E-03** | 3.90E-04 | **2.71E-03** |
| PANg | 4.55 | 3.15E-02 | **7.38E-02** | -4.033 | 2.01E-01 | 3.14E-01 | 8.506 | 4.51E-04 | **2.71E-03** | 1.82E-02 | **4.71E-02** |
| PANs | -0.1778 | 7.53E-01 | 7.75E-01 | -0.9 | 2.89E-01 | 3.85E-01 | 0.5353 | 4.10E-01 | 4.92E-01 | 5.68E-01 | 6.20E-01 |
| PANtotal | 12.95 | 7.37E-03 | **2.41E-02** | -7.383 | 3.01E-01 | 3.87E-01 | 21.634 | 8.87E-05 | **1.04E-03** | 6.33E-03 | **2.28E-02** |
| Course | 0.9683 | 3.28E-02 | 7.38E-02 | -0.9754 | 1.35E-01 | 2.31E-01 | 1.265 | 1.83E-02 | **4.71E-02** | 0.01121 | **3.36E-02** |

Table S5 Comparison of input clinical variables across NFBC *female*-subject subgroups

| Measures | Estimate | P value | Q value |
| --- | --- | --- | --- |
| WHR | -0.136587 | 1.32E-199 | **1.58E-198** |
| crp3dec | -1.2323 | 7.34E-06 | **8.01E-06** |
| FB_GLUK | -0.1977 | 1.27E-04 | **1.27E-04** |
| FS_INS | -5.5978 | 1.72E-51 | **5.16E-51** |
| FS_KOL | -0.51332 | 1.92E-09 | **2.56E-09** |
| FS_KOL_H | 0.17767 | 1.43E-10 | **2.15E-10** |
| FS_KOL_L | -0.3954 | 2.13E-07 | **2.56E-07** |
| FS_TRIGL | -0.8068 | 2.97E-31 | **7.13E-31** |
| HOMA_IR | -0.72122 | 1.27E-52 | **5.08E-52** |
| BMI | -6.1981 | 1.77E-100 | **1.06E-99** |
| SBP | -7.762 | 3.52E-13 | **6.03E-13** |
| DBP | -8.2871 | 6.93E-18 | **1.39E-17** |

Table S6 Comparison of input clinical variables across NFBC *male*-subject subgroups

| Measures | Estimate | P value | Q value |
| --- | --- | --- | --- |
| WHR | -0.284545 | 3.87E-224 | **4.64E-223** |
| crp3dec | -2.4734 | 1.45E-06 | **1.93E-06** |
| FB_GLUK | -0.19036 | 1.80E-03 | **1.80E-03** |
| FS_INS | -5.7274 | 4.17E-34 | **1.67E-33** |
| FS_KOL | -0.3737 | 9.20E-04 | **1.00E-03** |
| FS_KOL_H | 0.26044 | 1.60E-08 | **2.40E-08** |
| FS_KOL_L | -0.41459 | 2.91E-05 | **3.49E-05** |
| FS_TRIGL | -0.4324 | 1.26E-11 | **2.52E-11** |
| HOMA_IR | -0.72882 | 2.87E-33 | **8.61E-33** |
| BMI | -9.335 | 5.28E-59 | **3.17E-58** |
| SBP | -10.482 | 8.05E-12 | **1.93E-11** |
| DBP | -8.076 | 9.36E-10 | **1.60E-09** |
